# Spatial and Temporal Origin of The Third SARS-Cov-2 Outbreak in Taiwan

**DOI:** 10.1101/2022.07.04.498645

**Authors:** Jui-Hung Tai, Yu Ken Low, Selina Cai-Ling Wang, Hsin-Fu Lin, Tzi-Yuan Wang, Jann-Tay Wang, Yu-Shu Liu, You-Yu Lin, Charles S.P. Foster, Sebastiaan J. van Hal, Ya-Yun Lai, Shiou-Hwei Yeh, Sui-Yuan Chang, Pei-Jer Chen, Shu-Miaw Chaw

## Abstract

Since the first report of SARS-CoV-2 in December 2019, Taiwan has gone through three local outbreaks. Unlike the first two outbreaks, the spatial and temporal origin of the third outbreak (April 20 to November 5, 2021) is still unclear. We assembled and analyzed a data set of more than 6,000 SARS-CoV-2 genomes, including 300 from Taiwan and 5812 related sequences downloaded from GISAID as of 2021/12/08. We found that the third outbreak in Taiwan was caused by a single virus lineage belonging to Alpha (B.1.1.7) strain. This lineage, T-III (the third outbreak in Taiwan), carries a distinct genetic fingerprint, consisting of spike M1237I (S-M1237I) and three silent mutations, C5812T, C15895T, and T27869C. The T-III is closest to the sequences derived from Turkey on February 8, 2021. The estimated age of the most recent common ancestor (TMRCA) of T-III is March 23, 2021 (95% highest posterior density [HPD] February 24 - April 13, 2021), almost one month before the first three confirmed cases on April 20, 2021. The effective population size of the T-III showed approximately 20-fold increase after the onset of the outbreak and reached a plateau in early June 2021. Our results reconcile several unresolved observations, including the occurrence of two infection clusters at the same time without traceable connection and several airline pilots who were PCR negative but serum IgM-/IgG+ for SARS-CoV-2 in late April. Therefore, in contrast to the general notion that the third SARS-CoV-2 outbreak in Taiwan was sparked by two imported cases from USA on April 20, 2021, which, in turn, was caused by the partial relaxation of entry quarantine measures in early April 2021, our comprehensive analyses demonstrated that the outbreak was most likely originated from Europe in February 2021.

## Introduction

Since the first report of coronavirus disease 2019 (COVID-19) caused by *Severe acute respiratory syndrome coronavirus* 2 (SARS-CoV-2) in the December of 2019 in Wuhan, the virus has rapidly sparked an ongoing pandemic. SARS-CoV-2 is the third coronavirus causing severe respiratory illness in humans after SARS-CoV and Middle East respiratory syndrome coronavirus (MERS-CoV) (1-3).

After experiencing a series of SARS outbreaks in 2003 which caused 668 probable cases and 181 deaths (4), Taiwan has been exceedingly cautious of emerging disease and has strengthened its pandemic control measures. For example, the Central Epidemic Command Center (CECC) was established after the SARS epidemic in 2003, and was activated on 20 January 2020, before the first case of COVID-19 was identified in Taiwan. The control strategy implemented by the CECC was based on three essential components: border control, case identification and contact tracing, and containment.

As of February 2022, Taiwan has ended three local COVID-19 outbreaks, while the fourth is ongoing (Fig. 1). The first local outbreak was between January 28 and April 11, 2020 and involved 55 confirmed cases. Most of these local cases had a contact history or exposure to SARS-CoV-2 infected patients (5). The second local outbreak started on January 12 and ended on February 9, 2021. It was sparked by an intra-hospital infection and involved 21 cases. The third outbreak consists of two infection clusters and lasted for at least five months with more than 14,000 cases. Among them, all of the 614 cases tested by CECC are Alpha strain (B.1.1.7) of SARS-CoV-2 (6). The first cluster began with two airline flight crews (case 1078 and 1079) showing symptoms on April 17 and 18, respectively, after returning from the USA on April 16, 2021 (Table S1). They were diagnosed as COVID-19 positive on April 20, 2021 (7). On the same day, one pilot of the same airline tested positive for COVID-19 in Australia while on duty (8). This cluster was subsequently linked to staff working in a hotel in Northern Taiwan close to Taoyuan Airport where airline pilots and flight crews stayed during their quarantine (9) (Cluster I). The second cluster involved several local incidences in New Taipei City and Yilan County that later spread to many other counties (Cluster II). This cluster was first recognized on May 11, 2021, but a later survey found that the first case (case 1424) in this cluster showed symptoms as early as April 23, 2021 (10). Unlike the first two outbreaks, the spatial and temporal origin of this outbreak is still under debate.

**Figure 1.**
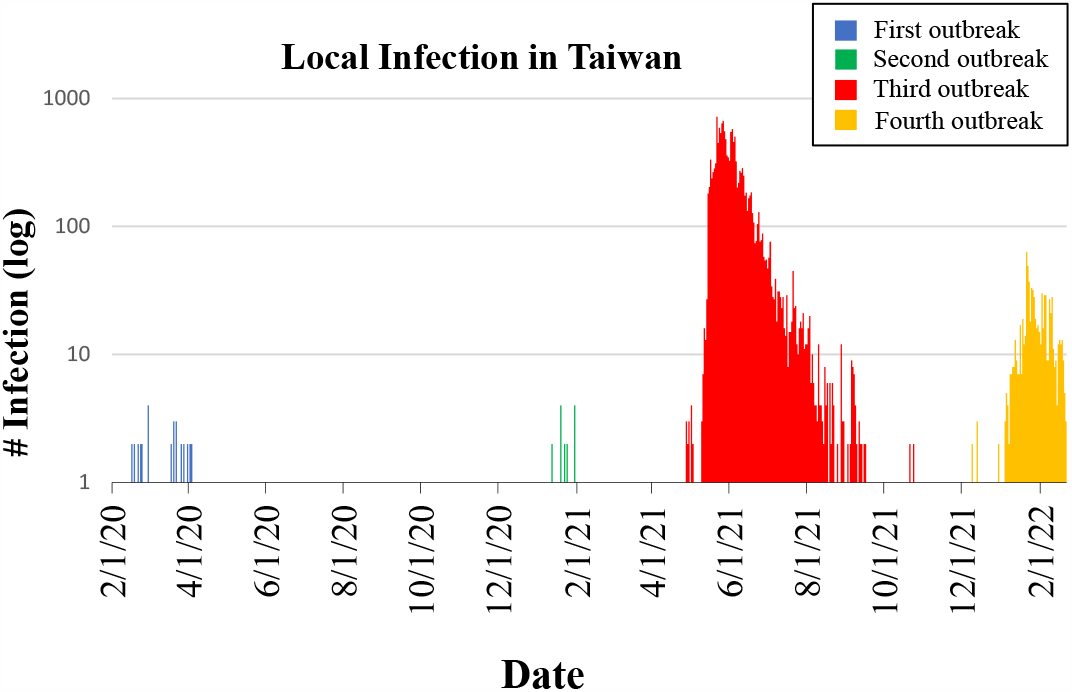
Number of local COVID-19 cases in Taiwan. Data were downloaded from COVID-19 DASHBOARD (https://covid-19.nchc.org.tw/) on February 22, 2022.

While it has been suggested that the third SARS-CoV-2 outbreak in Taiwan was sparked by two imported cases from USA on April 20, 2021 (11), which, in turn, was caused by the partial relaxation of entry quarantine measures in early April 2021 (12, 13), several unresolved observations remain to be answered. First, according to CECC, the dates of symptom onset in two infection clusters are very close (April 16 for the first cluster and April 23 for the second) (10). However, the relationships between the two clusters remain elusive. Second, several airline pilots and their family members were PCR negative but with serum IgM-/IgG+ in late April and early May 2021 (14, 15), suggesting they have probably been infected by SARS-CoV-2 earlier than April 20, 2021. While it is not entirely impossible that these cases were not linked to the third outbreak in Taiwan, the fact that there were only 1077 confirmed COVID-19 cases in Taiwan prior to April 20, 2021 (Fig. 1) makes it more probable that these cases are associated to the third outbreak.

Understanding the origin of the outbreak is important not only for epidemiological study but also for the design of future control measures. In order to clarify the origin(s) and interrelationship between the clusters, we sequenced and reconstructed a phylogeny of SARS-CoV-2 genomes. We find that the third outbreak is caused by a single virus lineage (T-III), descended from the Alpha variant of concern (B.1.1.7), which carries four distinctive mutations, including spike M1237I (S-M1237I) and three silent changes, from its most closely related sequences in Europe.

## Methods

### Sample preparation

Thirty-three specimens from SARS-CoV-2 positive patients were obtained (Table S1). One sample from Royal Prince Alfred Hospital, Australia was sequenced directly from extract using Nanopore (16). The remaining 32 specimens from individuals who were admitted to National Taiwan University Hospital (NTUH) were maintained in viral transport medium. Virus in the specimens was propagated in VeroE6 cells in Dulbecco’s modified Eagle’s medium (DMEM) supplemented with 2 μg/mL tosylsulfonyl phenylalanyl chloromethyl ketone (TPCK)-trypsin (Sigma-Aldrich, St. Louis, MI, USA). After one passage, culture supernatant was harvested when cytopathic effects (CPE) were observed in more than 70% of cells, and the culture supernatant was harvested for viral genomic sequencing. Viral RNA was extracted by QIAamp Viral RNA Mini Kit (Qiagen, Hilden, Germany) according to the instruction manual. In brief, 140 microliter of culture supernatant was used for RNA extraction and the extracted RNA was eluted using 40 μL RNase-free H2O. The RNA was stored at -20°C until use.

### Reverse transcription polymerase chain reaction (RT-PCR)

SARS-CoV-2 cDNA was generated from 100 ng of RNA in a RT-PCR reaction buffer containing 4 μL of 5X PrimeScript IV 1st strand cDNA Synthesis Mix (Takara Bio, Kusatsu, Shiga, Japan), 2 μL of 50 μM random hexamer primer, and variable amount of DEPC water to fill up to 20 μL of total reaction volume. Pre-heat at 30□ for 10 minutes, followed by 20 minutes of 42□, and then 15 minutes of 70□.

### Amplification of complete SARS-CoV-2 genomes with multiplex PCR

The “Midnight” amplicon protocol was used to generate 1200 bp amplicons. Briefly, PCR was used with 2·5 μL of cDNA product from RT-PCR in 22·5 μL buffer, containing 12·5 μL of Q5 Hot Start High-Fidelity 2X Master Mix (New England BioLabs, M0494S), 1·1 μL of 10 μM SARS-Cov2-Midnight-1200 primer (either Pool 1 or Pool 2)(17) (Integrated DNA Technologies, Coralville, IA USA), and 8·9 μL of nuclease-free water (Thermo Scientific, Waltham, MA, USA). Amplifications were performed with 30 seconds of 98□ for initial denaturation, followed by 25 cycles of 98□ for 15 seconds and 65□ for 5 minutes in a Veriti 96-Well Thermal Cycler machine (Applied Biosystems, Waltham, MA, USA). Each sample was separately amplified using both Pool 1 and Pool 2 primers. 20 μL PCR products were then purified with DNA Clean & Concentrator-5 (Zymo Research, Irvine, CA, USA). Amplicons were eluted with 25 μL of nuclease-free water (Thermo Scientific). DNA quality checks were done using a Nanodrop (Thermo Scientific) and 1·5% agarose gel electrophoresis (Fig. S1).

### Library preparation for Nanopore MinION sequencing

For each sample, 1 μL (0·5 μL from each pool of a same sample), approximately 50 ng, of purified PCR amplicons were used for library preparation. The KAPA HyperPrep Kit (Roche, Basel, Switzerland) was used in a 15 μL reaction for end repair and A-tailing. The reactions contained 1 μL of amplicon, 1·75 μL of End-repair & A-tailing Buffer, 0·75 μL of End-repair & A-tailing Enzyme, and 11·5 μL of nuclease-free water (Thermo Scientific). Reactions were done in 30 minutes at 20 □ and 30 minutes at 65 □. Each sample was then barcoded via ligation in a 27·5 μL reaction at 20□ for 15 minutes, with 2·5 μL of DNA Ligase, 7·5 μL of Ligation Buffer, 2·5 μL of Native Barcode (Oxford Nanopore Technologies, Oxford, UK), and 15 μL of A-tailed amplicon. After that, barcoded amplicons were purified with 33 μL (1.2X) of KAPA Pure Beads (Roche) by following the official protocol and eluted with 11 μL of nuclease-free water (Thermo Scientific, R0582). We then pooled 2·7 μL of each barcoded amplicon. The 110 μL of reaction for adapter ligation contained 65 μL of pooled barcoded amplicons, 5 μL of Adapter Mix II (Oxford Nanopore Technologies, EXP-NBD104), 30 μL of Ligation Buffer and 10 μL of DNA Ligase. After incubating in 20 □ for 15 minutes, libraries were purified with 110 μL (1X) of KAPA Pure Beads (Roche, 07983271001) and Short Fragment Buffer (Oxford Nanopore Technologies, SQK-LSK109) according to the official protocol and then eluted with 14 μL of Elution Buffer (Oxford Nanopore Technologies, SQK-LSK109).

### Nanopore MinION sequencing

We sequenced 220 ng of purified library with an Oxford Nanopore Technoogies MinION R.9.4.1 flowcell (FLO-MIN106) with the software MinKNOW Core version 19.12.5. Super-accurate basecalling was done using Guppy version 5.0.11 with default settings in GPU mode. The Nanopore reads of each sample were mapped to the reference genome Wuhan-hu-1 (EPI_ISL_402125) by *BWA-mem* [preprint] (18) to generate an alignment file. The average read depth is 992 per sample (Table S1). *BCFtools* (19) was applied to call variant (-i ‘QUAL > 20’) and generate consensus sequence.

### Data collection

A total of 267 complete and high coverage SARS-CoV-2 genomes from Taiwan with complete collection date were downloaded from the Global Initiative on Sharing Avian Influenza Data (GISAID, https://www.gisaid.org/)(20) on December 8, 2021. To find the SARS-CoV-2 genomes most closely related to the third outbreak in Taiwan, we used the Audacity Instant search tool in GISAID to search the database and used EPI_ISL_2455264 (case 1079) as the query. After excluding 42 sequences from Taiwan, 5,812 foreign sequences with fewer than or equal to 10 SNP differences were downloaded. All the sequences used in this study can be found in https://github.com/ala98412/T-III.

### M1237I frequency in different genetic backgrounds

We used tools within GISAID to provide a quick search of the database. We chose different SARS-CoV-2 strains in the drop-down menu “Variant” to get the count of sequences in each major variant strain (e.g., Alpha, Delta …), and further selected “Spike_M1237I” in the drop-down menu “Substitutions” to receive the count of sequences with the M1237I mutation in each genetic background. We calculated the frequency of M1237I in different genetic backgrounds via the count of sequences with the M1237I mutation divided by the total number of sequences representing each major variant.

### Sequence analysis and phylogenetic reconstruction

All sequences were aligned against the reference genome Wuhan-hu-1 (EPI_ISL_402125) by using MAFFT v7 (21). Nucleotide diversity, including number of segregating sites, Watterson’s estimator of θ (22), and nucleotide diversity (π)(23), was estimated using MEGA-X (24). Phylogenetic trees were also constructed by using the neighbor-joining method (25) based on Kimura’s two-parameter model in MEGA-X. The Nexus file for the haplotype network analysis was generated using DnaSP 6.0 (26) and input into PopART v1.7 (27) to construct the haplotype network using TCS software (28).

The times to the most recent common ancestor (TMRCA) of T-III lineage were estimated using an established Bayesian Markov chain Monte Carlo (MCMC) approach implemented in BEAST version 2.5 (29). The sampling dates were incorporated into TMRCA estimation. These analyses were performed using the Hasegawa-Kishino-Yano (HKY) model of nucleotide substitution assuming an uncorrelated lognormal relaxed molecular clock (30). We linked substitution rates for the first and second codon positions and allowed independent rates for the third codon position. We performed two independent runs with 1×10^7^ MCMC steps and combined the results. Log files were checked using Tracer (http://beast.bgio.ed.ac.uk/Tracer). Effective sample sizes were >300 for all parameters.

## Results

A total of 33 SARS-CoV-2 genomes were sequenced, including 32 from NTUH, Taiwan, and one from Royal Prince Alfred Hospital, Australia (Table S1). We also downloaded all available 267 genomes from Taiwan from GISAID as of 2021/12/08, to construct the phylogeny as shown in Fig. 2. Since the cases from the first outbreak had a contact history or exposures to different SARS-CoV-2 infected patients, they do not form a single cluster in the phylogeny. The sequences derived from the second local outbreak are presented in emerald green (Fig. 2). The third local outbreak consisted of the Alpha strain (B.1.1.7), divided into two main clusters, shown in Fig. 2b. All sequences in the basal lineage of Fig. 2b were from imported cases, whereas all 80 sequences in the more advanced lineage were local to Taiwan. Cluster I and II cases prior to May 10, 2021, are marked on the tree, and there is no clear differentiation between them. Subsequently, a steep rise in case numbers made it impossible to distinguish between the two clusters. Hereafter, we name this particular lineage as T-III, which stands for the third outbreak in Taiwan, and note that the T-III belongs to Pangolin lineage B.1.1.7. The nucleotide changes among T-III are listed in Table S2.

**Figure 2.**
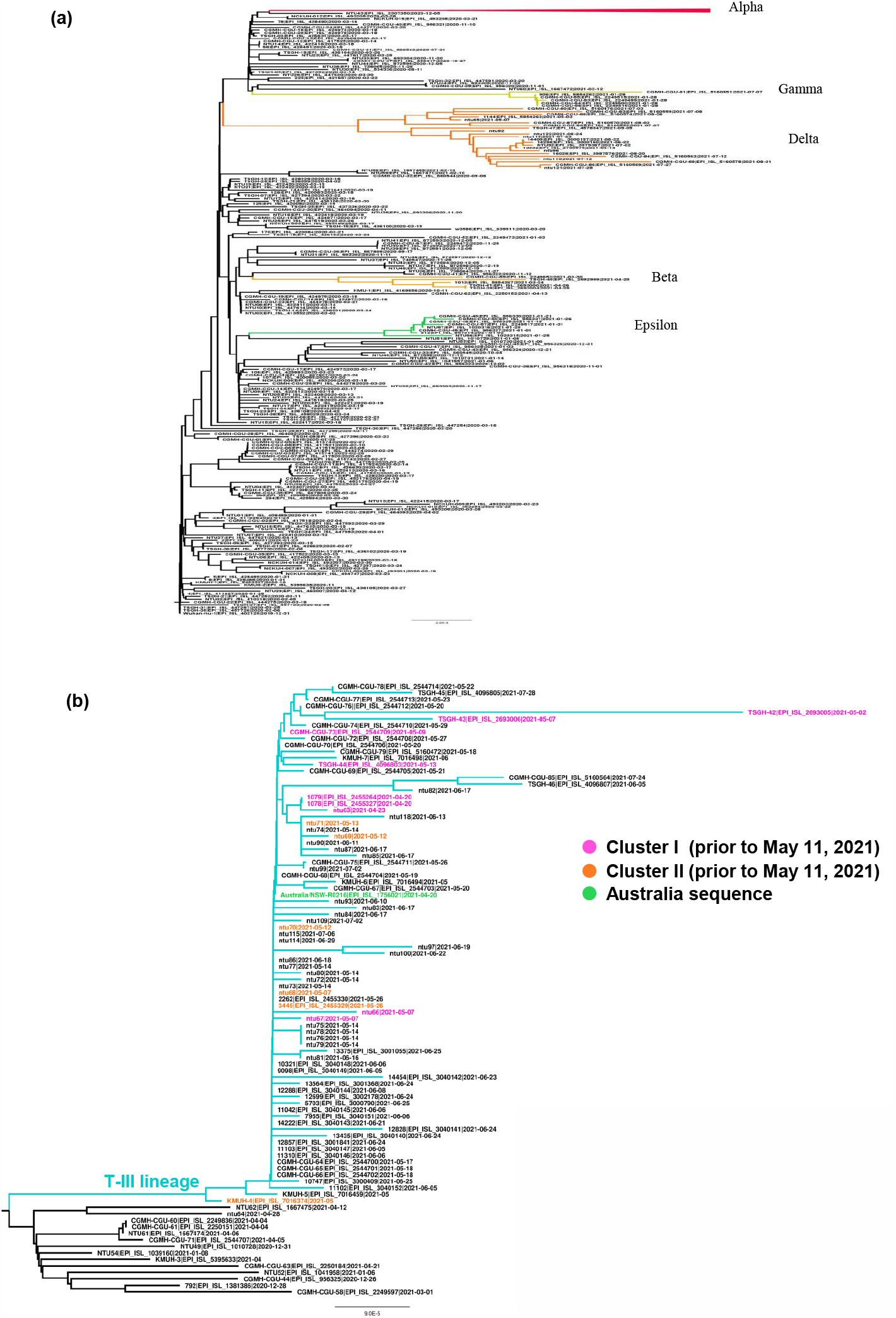
Phylogeny of (A) all SARS-CoV-2 and (B) Alpha strain only genomes from Taiwan. In (B) the black branches are imported, and colored branches are local cases. Australia sequence was colored in green. Cluster I and cluster II, both of which were collected prior to May 11, 2021, are colored in purple and orange, respectively.

### Spatial and temporal origin of the third local outbreak in Taiwan

In order to search for the spatial origin of T-III, the sequence recovered from the earliest case in the third outbreak (case 1079: EPI_ISL_2455264, Table S1) was used to search against the GISAID database. There were 5,812 sequences with ≤ 10 nucleotide differences from EPI_ISL_2455264 as of November 21, 2021. Phylogenetic reconstruction including all 5,812 sequences and T-III demonstrate that the latter is a distinctive lineage (Fig. S2). The vast majority of sequences closely related to T-III were from Europe, including Turkey (Fig. S2b). We also tested whether the sequence from Australia is unique to T-III. All available Alpha strain genomes from Australia collected during April 2021 were downloaded from GISAID as of February 17, 2023 and compared with T-III (Fig S3). The sole sequence (EPI_ISL_1756021) belonging to T-III from Australia is distinct from the other Alpha strains collected at the same time. As EPI_ISL_1756021 was from a flight crew from Taiwan, the result supports the notion that the T-III is unique to the third outbreak in Taiwan.

Haplotype network analyses of the T-III lineage reveals 48 haplotypes. The network shows that T-III differs from the outgroups in four mutation steps (Fig. 3), including two synonymous mutations, C5812T and C15895T, in Orf1ab, one nonsynonymous mutation G25273C (M1237I) in Spike, and one T27869C mutation in a non-coding region (Table S3). The closest outgroup haplotype consists of four sequences, including two sequences from Turkey that were collected on February 8, 2021 (EPI_ISL_1097034, 1097035). The rest were collected after the onset of the third outbreak in Taiwan (Table S4). Further database mining confirmed that of 1,140,328 Alpha strain genomes examined as of December 11, 2021, only the lineage T-III possessed the four above mentioned mutations, which form a distinctive genetic fingerprint. Within T-III, the network forms a star-like shape centered on a core haplotype comprising 26 sequences. Most of the remaining haplotypes are directly connected to this major haplotype. Of the three cases identified on the first day of the third outbreak (April 20, 2020), the case from Australia belongs to the major haplotype, with the rest (1078 and 1079) one mutation away.

**Figure 3.**
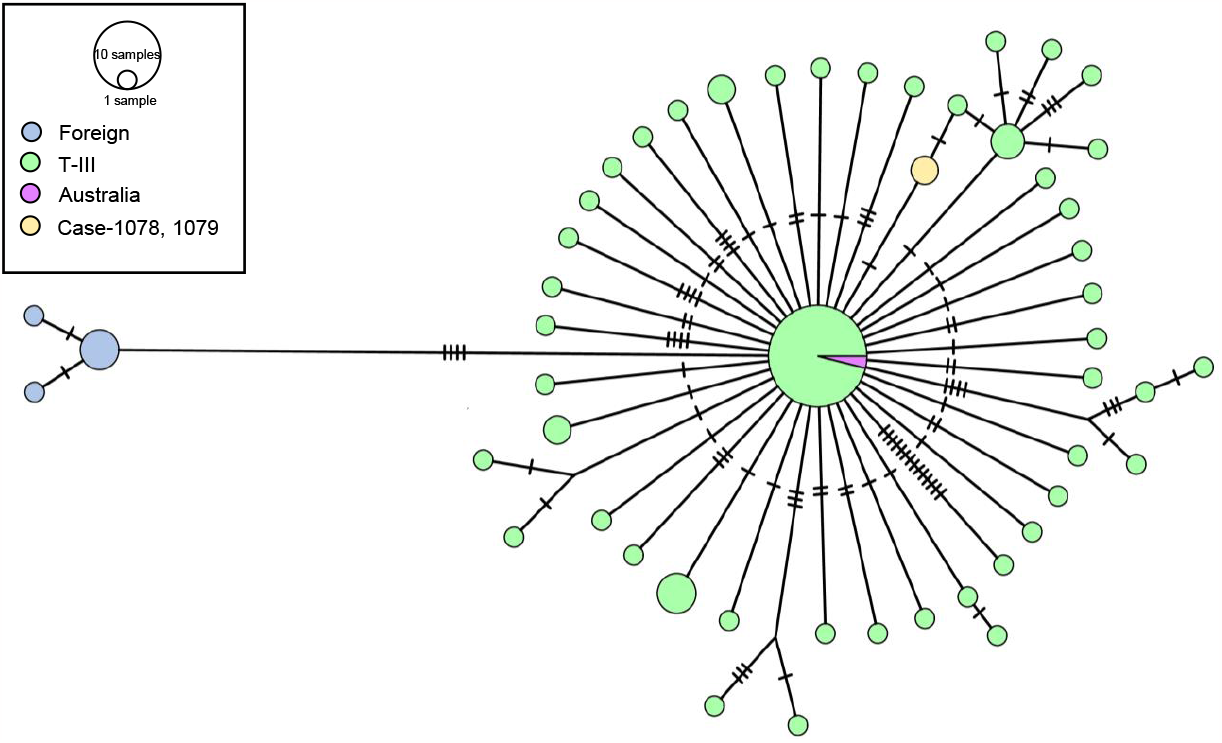
Haplotype network of SARS-CoV-2 genomes from the third local outbreak. The haplotype network was constructed by the median joining algorithm. Circle areas are proportional to the number of sequences. Mutational steps between haplotypes are symbolized by dashes. Haplotypes in yellow and pink are from Airline flight crews, who were the first three cases during the third outbreak. Cases 1078 and 1079 (in yellow) were diagnosed as COVID-19 positive on April 20 (CECC 2021b). On the same day, one pilot of the same airline tested positive for COVID-19 in Australia (in pink) while on duty (CECC 2021f).

The estimated substitution rate of T-III is 9.8×10^−4^ substitution per site per year (95% highest posterior density [HPD] 5.6×10^−4^ – 1.4×10^−3^) which is close to 8.4 × 10^−4^ /site/year of SARS-CoV-2 (31), and 7.5□×□10^−4^ /site/year of strain B.1.1 (32). The results also suggest that a single passage in Vero E6 cells would not significantly influence the number of mutations on SARS-CoV-2 sequences. The date of the most recent common ancestor (TMRCA) of T-III is in late March (3/23/2021; 95% HPD February 24 - April 13, 2021) (Fig. 4), almost one month before the first three confirmed cases on April 20, 2021. We noticed that the mutation rate of T-III during this epidemic was almost constant (Fig. S4). Effective population size of the T-III lineage increased approximately 20-fold after the onset of the outbreak and reached a plateau in early June. The estimated demographic expansion of T-III is consistent with epidemiological data (Fig. 1). We noticed that demography in Fig. 4 does not capture the population decline after July 2021 as shown in Fig. 1. This finding is because most sequences used in this analysis were collected before June 2021 (Table S1), with only four sequences obtained after July 2, 2021. As we are interested in the outbreak origin, this sampling strategy should not affect our conclusions.

**Figure 4.**
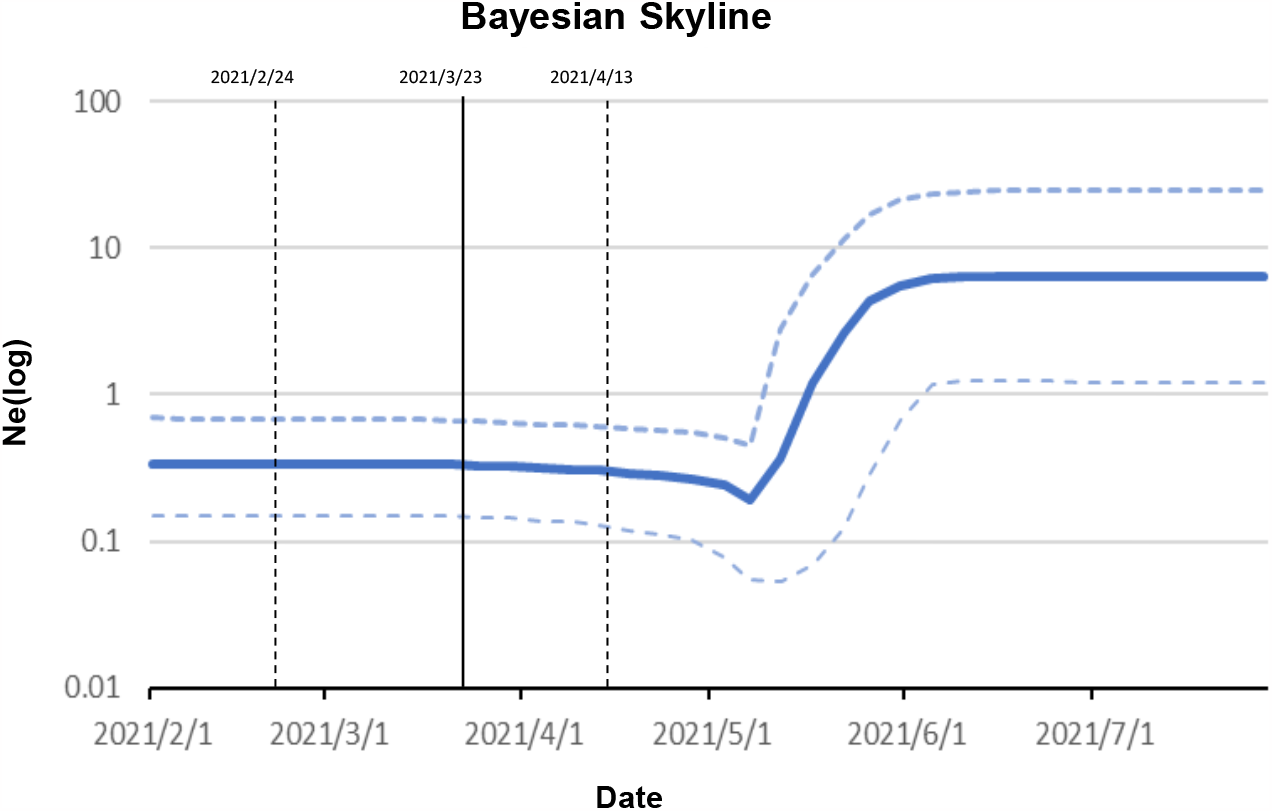
The epidemic growth curve of SARS-CoV-2 genomes from the third local outbreak. The three lines are the median (blue line) and 95% HPD intervals (dashed lines) of the Bayesian skyline plot (m = 5). Vertical solid line indicates the estimated time to the most recent common ancestor with 95% HPD intervals in dashed lines.

Rapid population expansion can also be revealed by contrasting patterns of genetic variation estimated using different approaches. The Watterson’s estimator of θ (6·95 × 10^−4^) is approximately seven times higher than the nucleotide diversity (π) (1·05 × 10^−4^), leading to significantly negative Tajima’s D (−2·82, p < 0·001) (Table S5). Because θ is strongly influenced by rare mutations, which are common during recent population expansion or after positive selection (33), it is a better estimator of genetic diversity for T-III.

## Discussion

Our results provide solid evidence that the third local outbreak in Taiwan was caused by a single lineage, T-III. This does not in itself mean that two clusters of infections (see Introduction) have a single common origin. For example, among 293,742 sequences analyzed during the first year of the SARS-CoV-2 pandemic, the most abundant haplotype was sampled 3,466 times from across 53 countries (Table S6). It is possible that the haplotype exhibited some transmission advantage, making it widespread. Under this scenario, two clusters of infection may be caused by the same lineage imported from different sources. However, as the T-III lineage bearing a combination of four unique mutations is exclusive to Taiwan, it seems highly unlikely that this lineage was imported from different sources.

Our estimation that T-III originated from Europe with TMRCA on March 23, 2021 (95% HPD February 24, 2021 □ April 13, 2021) reconciles several unresolved observations. First, the first two cases (case 1078 and 1079) of the outbreak shared identical sequences, indicating that they were from the same source. Nevertheless, the cycle threshold (Ct) values at the time of diagnosis (April 20, 2021) were 29 and 17 for case 1078 and 1079, respectively, suggesting that they were infected at different times (7). Second, several airline pilots and their family members were PCR negative but with serum IgM-/IgG+ in late April and early May 2021 (14, 15). It has been shown that IgM levels increase during the first week after a SARS-CoV-2 infection, peak after two weeks, and then recede to near-background levels in most patients. IgG is detectable one week after disease onset and is maintained at a high level for a long period (34). Consequently, IgM negative but IgG positive individuals have probably been infected by SARS-CoV-2 earlier than April 20, 2021. Third, according to CECC, the dates of symptom onset in two seemingly unrelated infection clusters are very close (April 16 for the first cluster and April 23 for the second) (10). As our phylogenetic analysis reveals that all sequences in the third outbreak have a single origin, the occurrence of two infection clusters at similar time without traceable connection demonstrates that the virus may have been cryptically circulating in the community undetected. Consequently, the origin of the third outbreak was most likely prior to April 20, 2021.

As most sequences closely related to T-III were from Europe, our results disagree with the notion that the outbreak was imported by cases 1078 and 1079 from the USA (11). Among four sequences closest to T-III, only two from Turkey were collected before April 20, 2021 (on February 8, 2021). The rest appeared after April 20 and cannot be associated with the third outbreak. Consequently, the lineage leading to the third outbreak was most likely introduced from Europe, perhaps Turkey, by infected travelers in February 2021 (Fig. 3).

There are several possible routes by which SARS-CoV-2 can be introduced by incoming travelers. First, some carriers may have the virus undetected during quarantine. Although the mean incubation period ranges from five to seven days, it can be longer than 14 days (35, 36). Indeed, there has been much debate about whether changing from the mandatory five day home quarantine plus nine day autonomous health management (or the so-called “5+9” strategy) to three day home quarantine plus 11 day autonomous health management (“3+11”) for flight crews on April 15, 2021 (37) was the cause of the third outbreak (12, 13). Since the outbreak likely originated in mid-March, as our analyses demonstrate, the possibility that changing strategies from “5+9” to “3+11” directly caused the outbreak can be ruled out.

Second, there may have transmission from people associated with quarantine hotels to the community. For example, from December 2021 to March 2022 alone, 15 infection clusters occurred in quarantine hotels (38). Since one of the two infection clusters in the third outbreak directly link to a hotel where many flight crews stayed during their quarantine (see Introduction), it is most likely that the outbreak was introduced from this hotel during late February to early April 2021. The virus was then undetected while spreading within the community until late April 2021.

Our results demonstrate that even a small number of imported cases can undermine the strict control measures in Taiwan (39). We also show that phylogenetic approaches can be used to trace the outbreak derived from local spread by infected travelers (40). Although Taiwan has been very cautious of emerging disease and has strengthened its control measures, the SARS-CoV-2 genomes have not been sampled in proportion to the actual size of infection and made publicly available in a timely fashion. This situation severely hinders efforts to trace the underlying transmission patterns of spread. Consequently, policies that include real-time public-sharing and transparency for infectious disease surveillance and control are critically and urgently needed. Blackouts in information sharing and transparency can cause information disparity and suspicion, which in turn hamper pandemic control efforts (41).

## Availability of Data and Materials

The National Taiwan University Hospital, Taiwan and the Royal Prince Alfred Hospital, Australia sequences have uploaded to Global Initiative on Sharing Avian Influenza Data (GISAID, https://www.gisaid.org/). The remaining genome sequences were downloaded from GISAID as well. All the sequences used in this study can be downloaded from https://github.com/ala98412/T-III.

## Authors’ Contributions

JHT, SCLW, and HFL analyzed the data. SYC and SHY guided the analyses. JHT, YKL, SCLW, PJC, HYW, CSPF drafted and revised the manuscript. YKL, Ya-Yun Lin, You-Yu Lin, JTW, YSL, and SYC prepared samples. TYW, CSPF, SJH, and You-Yu Lin sequenced samples. PJC, SMC, and HYW designed the study, obtained funding, and wrote the manuscript.

## Acknowledgement

The authors thank four anonymous reviewers for constructive comments. We also thank Prof. Chwan-Chuen King for her comments on the early version of this manuscript.

