## Supplement Figures for "Spatial and Temporal Origin of The Third SARS-Cov-2 Outbreak in Taiwan"

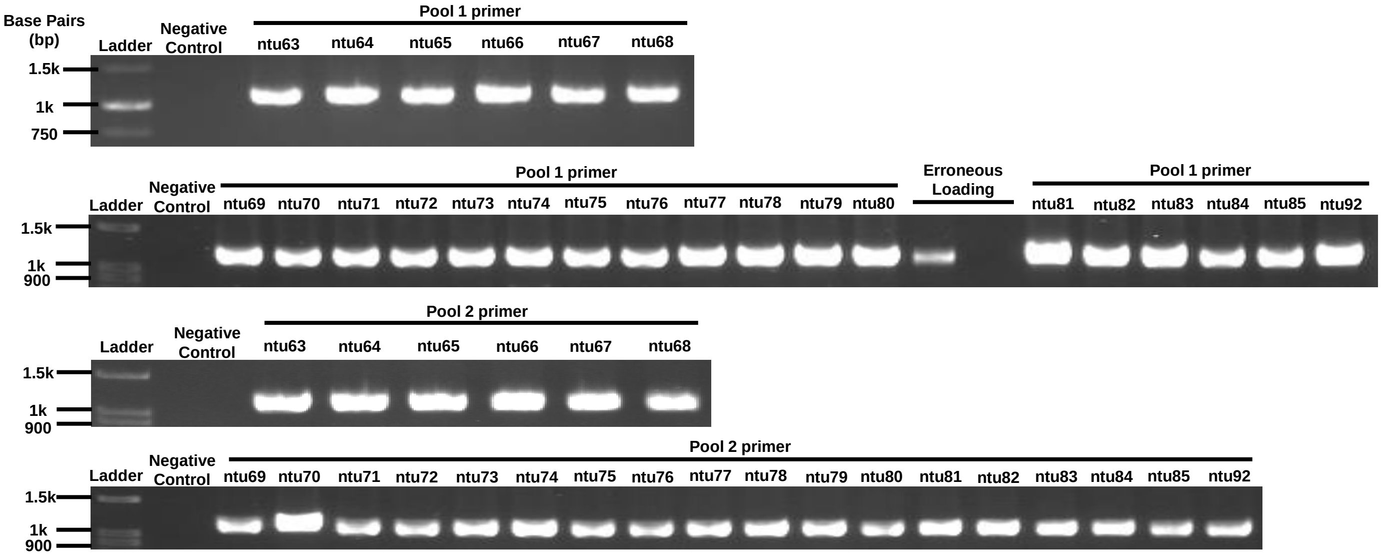


**Fig. S1 Agarose gel electrophoresis of amplicons.** The representative gel picture of 1,200 bp amplicons generated from SARS-CoV-2 cDNA using the “Midnight” amplicon multiplex PCR (see Methods for more details).

**(a)**

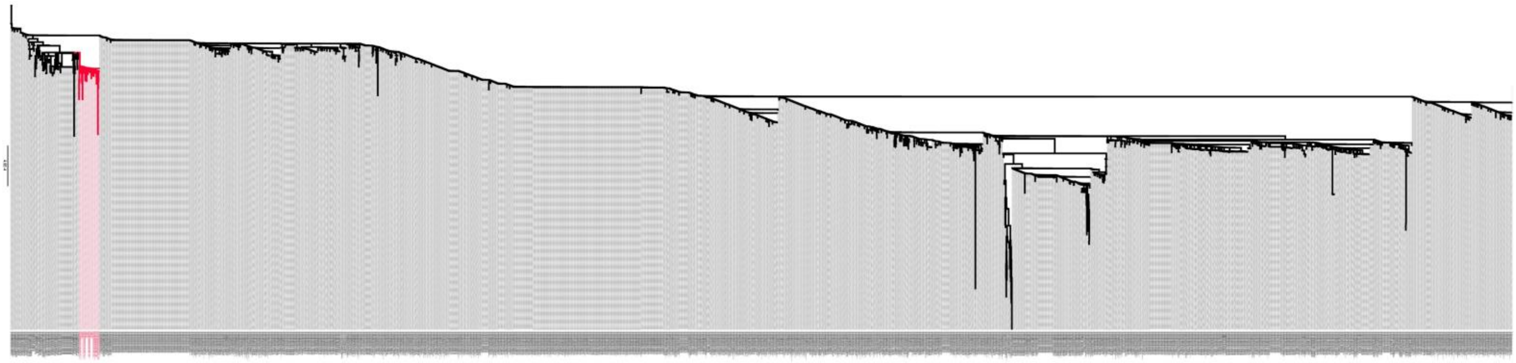

**(b)**

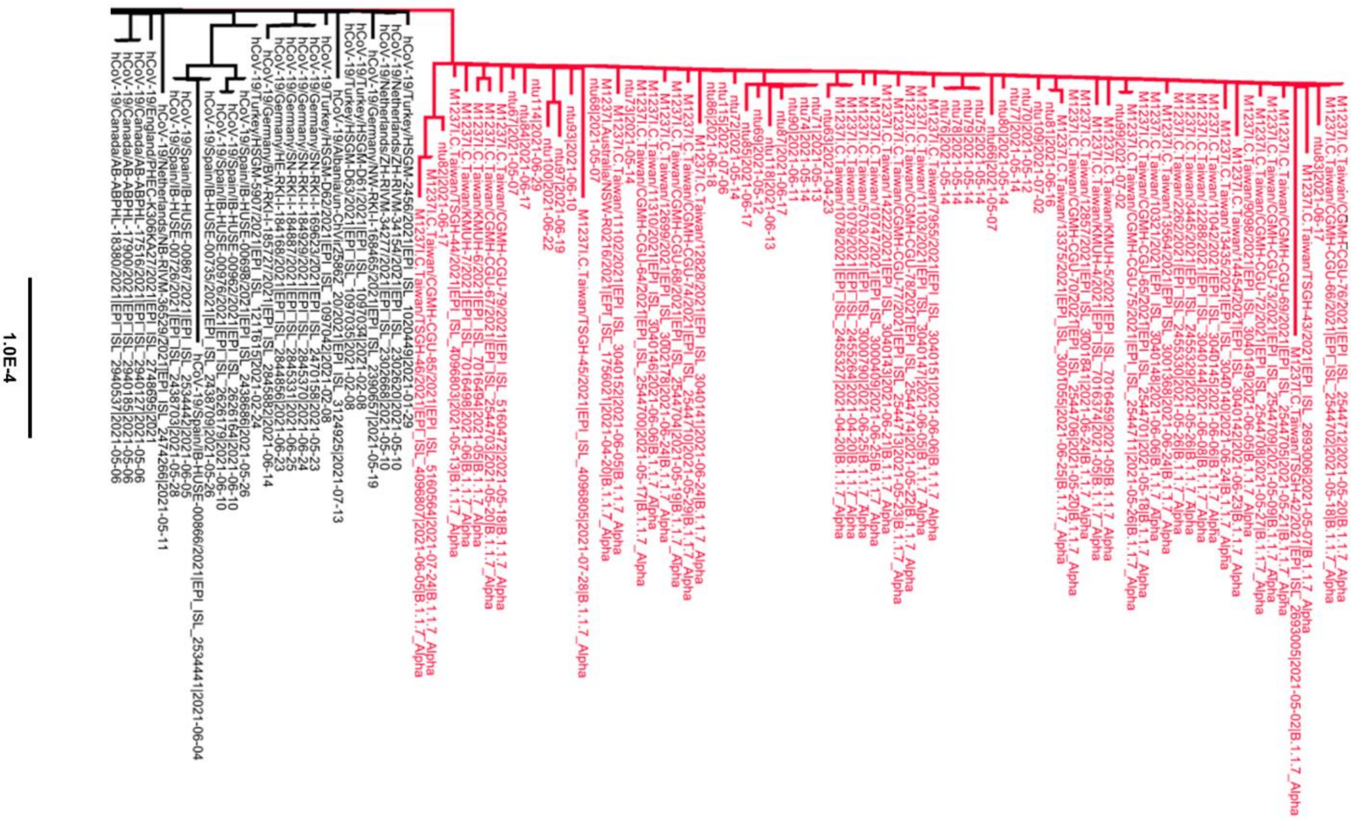


**Fig. S2 Phylogenetic tree of T-III lineage and their 5812 close-relative sequences.** Branch colored in red is T-III lineage. (a) The entire phylogenetic tree. (b) A closer look for T-III lineage.

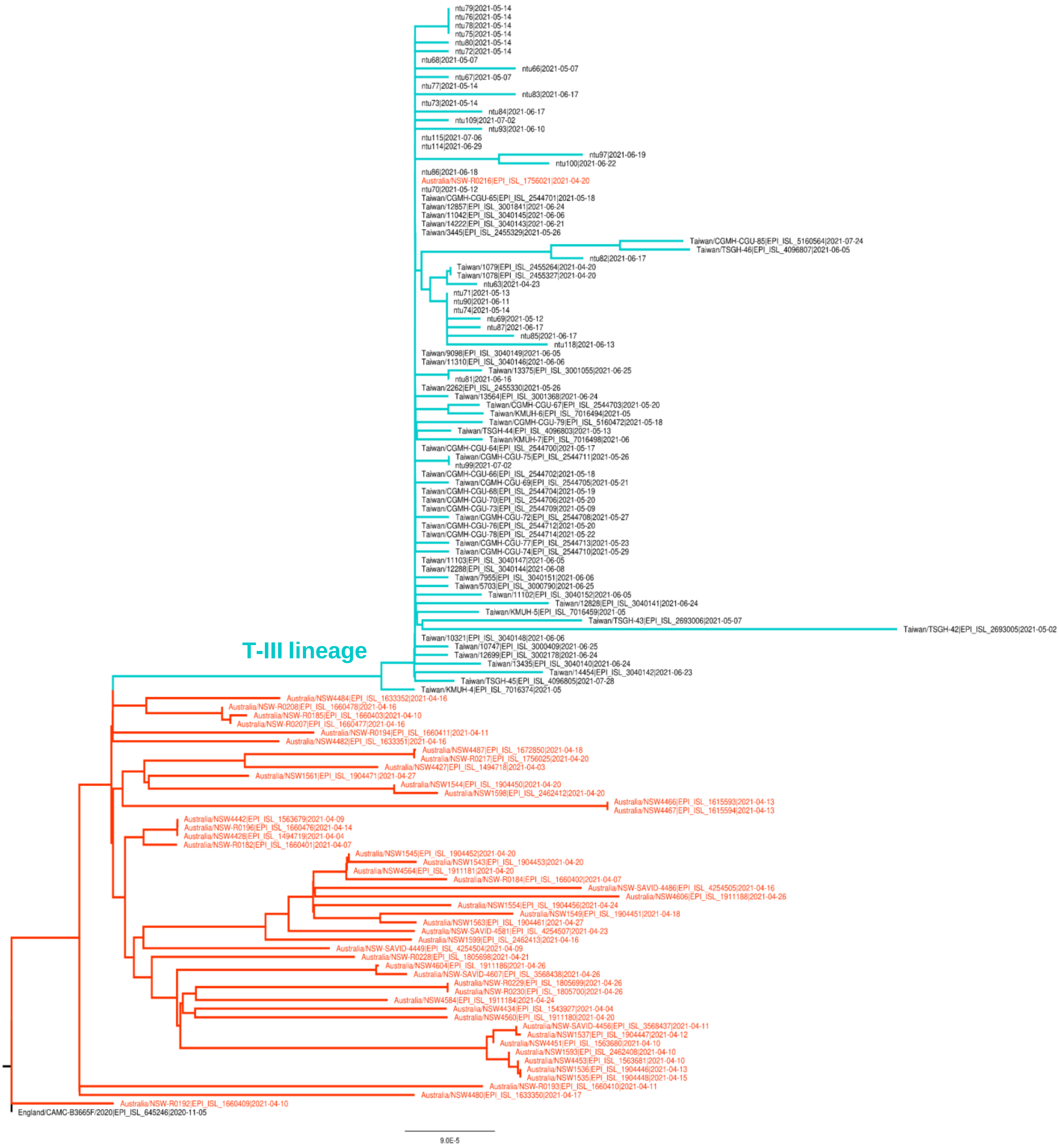


**Fig. S3 Phylogenetic tree of T-III lineage and alpha strain sequences from Australia collected during April, 2021.** Branches in orange are from Australia, whereas in cyan are from Taiwan. Brach leading to T-III lineage is marked.

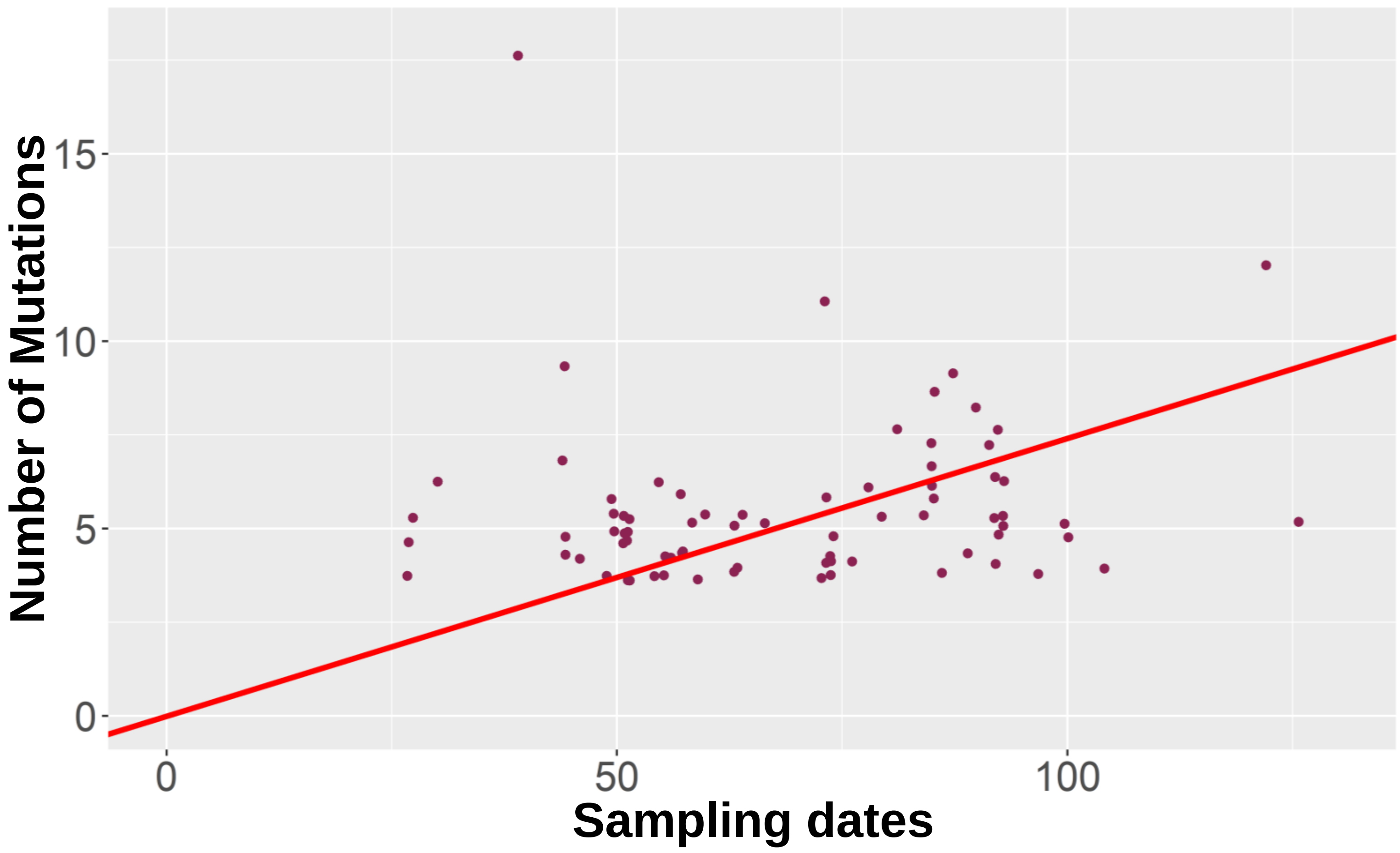


**Fig. S4 Number of mutations versus sampling dates.** The EPI_ISL_1097034 from Turkey was used as the outgroup (see Fig. 3 and Table S3 for details). Each dot represent a sequence of T-III. The number of mutations is the difference between sequences of T-III and EPI_ISL_1097034. The sampling dates are measured as the number of days away from March, 23 2021, the date of the most recent common ancestor estimated by an established Bayesian Markov chain Monte Carlo (MCMC) approach implemented in BEAST (27). A simple linear regression model (red line) was used to show the linearity of the molecular clock ($r^2 = 0.7623$ and p-value < 2e-16).
